# Identification of novel small molecule chaperone activators for neurodegenerative disease treatment

**DOI:** 10.1101/2024.07.17.603883

**Authors:** Anita Ho, Fiona Jeganathan, Magda Bictash, Han-Jou Chen

## Abstract

A pathological hallmark of neurodegenerative disease is the accumulation of aberrant protein aggregates which contribute to the cytotoxicity and are therefore a target for new therapy development. One key machinery to manage cellular protein homeostasis is the chaperones proteins, the heat shock proteins (HSPs) which are known to target aberrant proteins including TDP-43, tau and amyloid and rescue neurodegeneration in various disease models. As an attempt to target HSP activation for neurodegeneration therapy, we develop a drug screening assay to search compounds that activate the master regulator of HSPs, the transcription factor heat shock factor 1 (HSF1). As HSF1 is bound by HSP90 which prevents its activation, we develop a NanoBRET assay, which allows us to monitor and quantify the HSF1-HSP90 interaction in living cells to screen for compounds disrupting this interaction and thereby releasing HSF1 for activation. After the assay was optimised and validated for its robustness and reliability, a two thousand compound library was screened which produced 10 hits including a couple of known HSP90 inhibitors. Follow-up functional study showed that one of the hit oxyphenbutazone (OPB) significantly reduces the accumulation of insoluble TDP-43 in a cell model, displaying no signs of stress or toxicity. Further works are needed to characterise the mode-of-action of OPB in HSF1 activation and aberrant protein reduction, and the physiological impact of OPB on TDP-43 needs to be examined in an in vivo model. However, this study demonstrates a viable strategy for new drug discovery in targeting aberrant proteins and identifies potential candidates for translation into neurodegenerative disease treatment.

## Introduction

Neurodegenerative disorders (NDs) are characterised by the progressive loss of neurons. Depending on the specific brain region affected, the outcome would range from issues related to movement dysfunction to the development of dementia. They affected an estimate of 57.4 million of people worldwide in 2019, with a significantly increased related burden in recent years causing a significant sociological strain on our growing ageing population (Collaborators, 2022). Despite many recent developments and newly approved treatments, currently, there is still no cure for any ND. Some NDs have a familial component with known associated genetic mutations or genetic risks, many are sporadic where environmental risks and other yet-to-identified factors contribute to the disease development. Even with the heterogeneity of disease causes and sites affected, most NDs share a common pathological hallmark of the accumulation of misfolding aberrant protein aggregates in the affected neuronal areas. The widely recognised ND associated protein aggregates include amyloid-β and tau in Alzheimer’s disease (Allsop, 2000; Alzheimer et al., 1995), α-synuclein in Parkinson’s disease (Baba et al., 1998; Spillantini et al., 1998; Spillantini et al., 1997), and TDP-43 in amyotrophic lateral sclerosis (ALS) and frontotemporal lobar degeneration (FTD) (Arai et al., 2006; Neumann et al., 2006). Studies in various cell and animal models demonstrate that the accumulation of these aberrant protein aggregates leads to neuronal cytotoxicity, eventually neurodegeneration and neuronal death (Calabresi et al., 2023; Hayes & Kalab, 2022; Sehar et al., 2022; Virgilio et al., 2022). Therefore, the targeted clearance of protein aggregates is a key strategy for therapeutic intervention.

In healthy cells, protein synthesis, folding and degradation are tightly controlled and maintained by the chaperone system. The main players of the chaperone system are the heat shock proteins (HSPs) and the master transcription factor, heat shock factor 1 (HSF1). Chaperone proteins such as HSPs are controlled by HSF1 and undertake a wide range of cellular functions including protein homeostasis, cell survival and cell metabolism (Binder & Pedley, 2023; Kim et al., 2013; Margulis et al., 2020). In unstressed condition, some HSPs are present in steady level to facilitate the daily cellular function and maintain homeostasis. During stress, HSPs are further induced through a mechanism called the heat shock response (HSR) when HSF1 is released from its inhibitory binding from HSP90 (Zou et al., 1998), trimerizing and binding to the heat shock element (HSE) in the promoter region of HSPs and promoting the expressions of HSPs (Kingston et al., 1987; Sarge et al., 1993; Westwood et al., 1991). Timely and effective activation of HSR enhances the capacity of the cells to resolve accumulated unfolded/misfolded protein and to survive and recover from the stress (Kurop et al., 2021).

Interestingly, reduced HSR efficiency is observed in aged cells with a further down-regulation of key chaperone proteins observed in various neurodegenerative disorders (Brehme et al., 2014; Chen et al., 2016; Margulis et al., 2020). Reduced HSF1 protein levels are also found in AD and ALS mouse models as well as in ALS patients (Chen et al., 2016; Trivedi & Jurivich, 2020). Together, these studies suggest that the compromised chaperone system may play a role in the build-up of aberrant proteinopathies that eventually lead to neurodegeneration. Indeed, studies by us and other scientists have demonstrated that up-regulating key HSPs can reduce the accumulation of protein aggregates and enhance the cell survival rate (Ash et al., 2010; Barmada et al., 2010; Chen et al., 2016; Stallings et al., 2010; Wils et al., 2010). For example, HSP70 and DNAJA2 have been shown to inhibit the nucleation and elongation of tau, preventing the sequestration of tau into tau aggregates (Kundel et al., 2018; Mok et al., 2018); DNAJB6 interacts with amyloid-β and inhibits the formation of amyloid nuclei (Osterlund et al., 2020); HSP70 and DNAJB2a facilitate the refolding of insoluble TDP-43 and enhance cell survival (Chen et al., 2016); and overexpression of DNAJB2a in SOD1 transgenic mice reduced SOD1 aggregation (Novoselov et al., 2013). These findings suggest a potential treatment via the manipulation of the HSR pathway.

To date, the majority of attempts to activate this pathway are via HSP90 inhibition to release HSF1 and these have shown some neuroprotective effects in cellular and animal models (Chen et al., 2014; Fujikake et al., 2008; Waza et al., 2005). However, HSP90 is an abundant and important chaperone with many important client proteins that are involved in the cell cycle and cell survival pathways (Taipale et al., 2010), the inhibition of which would inevitably repress the HSP90-mediated survival signalling leading to cytotoxicity, therefore unsuitable for neurodegeneration therapy (Aguila et al., 2014; Garg et al., 2016; Sanchez et al., 2020). This has led us to explore other compounds that could specifically disrupt interactions between HSP90 and HSF1 to activate the HSR as a potential therapy.

To search for new HSR activators via disruption of HSF1 and HSP90 interaction, we established a live cell-based reporting assay utilising a NanoBRET system to monitor the interaction of HSF1 and HSP90. When optimised and validated, this assay was used in a compound library screen to test ∼2200 compounds from a bioreference library for inhibitors of HSF1-HSP90 interaction. A selection of hit compounds was followed up, one of which demonstrated a reduction in insoluble TDP-43 proteins and has the potential for further therapeutic development.

## Methods and material

### Plasmids, antibodies and reagents

NanoBRET expression plasmids including the p53 and MDM2 positive control were purchased from Promega (NanoBRET PPI Starter Kit, N1811, Promega). HSP90 and HSF1 cDNA were cloned into the NanoBRET expression plasmid to generate the Nluc- and HT-tagged HSP90 and HSF1 constructs. The GFP-TDP-43 in pEGFP-C1 plasmid was generated and used in previous studies (Chen et al., 2016; Nishimura et al., 2010).

Primary antibodies used in this study included: mouse anti-Actin (1:5000, A1978, Sigma), rabbit anti-GAPDH (1:1000, #2118S, Cell Signaling), rabbit anti-HSF1 (1:50,000, ab52757, Abcam), mouse anti-HSP70 (1:1000, ab5439, Abcam), rabbit anti-HSP90 (1:10,000, ab203126, Abcam), rabbit anti-TDP-43 (1:2000, 10782-2-AP, Proteintech) for immunoblotting, mouse anti-phospho TDP-43 (1:2000, Cosmo Bio), rabbit anti-PABP (1:1000, ab21060, Abcam) for immunofluorescence staining.

Secondary antibodies used in this study included: goat anti-Mouse IgG (H+L) Secondary Antibody, DyLight 800 4X PEG (1:10000, Thermo Scientific), goat anti-Rabbit IgG (H+L) Secondary Antibody, DyLight 680 (1:10000, Thermo Scientific) for western blot analysis, goat anti-Mouse IgG (H+L) Secondary Antibody, DyLight 550 (1:500, Thermo Scientific), goat anti-Rabbit IgG (H+L) Secondary Antibody, Alexa Fluor 647 (1:500, Thermo Scientific) for immunofluorescence staining.

Compound and reagents used in this study included: 17-AAG (1515/1, Bio-Techne & A8476, Sigma), Arsenite (106277, Sigma), BAI1 (S8865, Selleck Chemicals LLC), DMSO (D2438, Sigma), Oxyphenbutazone (OPB) (SML0540, Sigma), XIB4035 (SML1159, Sigma).

### Cell Culture and DNA transfection

HEK293T cells were cultured using Dulbecco’s modified Eagle medium (DMEM), high glucose, GlutaMAX Supplement (10566016, Gibco) supplemented with 10% of foetal bovine serum (FBS), and maintained at 37°C, 5% CO_2_. Cells were plated a day before transfection and media were refreshed before plasmid DNA transfection using Lipofectamine™ 2000 Transfection Reagent (#11668019, Life Technologies/Thermo Fisher Scientific) per manufacturer’s instructions. Cells were left for 48 hours after transfection to be harvested for analysis unless otherwise stated.

### Compound plates for screening

The 2100 Enamine Bioreference compound library (manufacture source, provided by ARUK-DDI at UCL) was used for this study. 10 µM of the compound library diluted in DMSO were stamped in 96-well white-walled tissue culture plates (#3917, Costar) with vehicle (DMSO) and positive (17-AAG, 10 µM) control (as shown in Supplementary Figure 3A), and stored at −20°C prior to the screen. Plates were defrosted at room temperature for 30 minutes before the addition of HaloTag/Nluc transfected cells as described below.

### NanoBRET assay

The NanoBRET assay was adapted from the technical manual provided in the Promega NanoBRET Protein:Protein Interaction System (N1662, Promega). HEK293T cells were plated in a 6-well plate, or a 10 cm dish the day before transfection. On the day of transfection, cells were transfected using Lipofectamine 2000 Transfection Reagent, with a donor-to-acceptor ratio of 1:150 (i.e. 100 ng of NanoLuc (Nluc) Luciferase construct and 15 μg of HaloTag construct per 10 cm dish) unless otherwise stated. 24 hours after transfection, cells were trypsinised and resuspended in phenol red-free Opt-MEM (#11058021, Thermo Fisher Scientific) containing 5% FBS at a concentration of 5×10^5^ cells per ml mixing with HaloTag 618 ligand (1 µl/ ml). 100 µl of the cell mixture was plated per well of the 96-well plate containing the screened compound. The plates were then incubated for 24 hours at 37°C before the NanoBRET assay.

To measure the NanoBRET signal, Nano-Glo substrate was diluted in phenol red-free Opti-MEM at the concentration of 10 µl/ml and 25 µl of the substrate mixture was added to each well, using the injection system of CLARIOstar plate reader (BMG Labtech), following with a 30 second double orbital shaking at 300 rpm. Donor (450-80nm) and acceptor (610-100nm) emissions were immediately measured after substrate addition using CLARIOstar plate reader.

### Data and statistical analysis

The NanoBRET signal (mBU ratio) was calculated using the following formula:

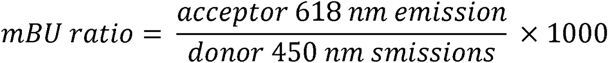

When optimising the NanoBRET assay for HSP90-HSF1 interaction, the corrected mBU ratio was used to determine the specificity of the assay, which factors in the donor-contributed background or bleed-through. The corrected mBU ratio is calculated by subtracting the mBU ratio of the experimental samples with HaloTag ligands from the mBU ratio without HaloTag ligands:

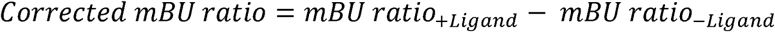

The fold change between with and without HaloTag ligands as well as between DMSO and 17-AAG controls were calculated to examine the separation using the following equation:

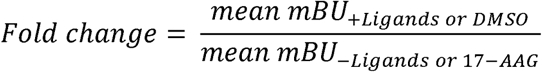

Z factor and coefficient of variation (CV) were used to measure the overall assay quality and robustness (Zhang et al., 1999). Z factor between with ligand and without ligand as well as between the DMSO and 17-AAG controls is calculated using the following equation:

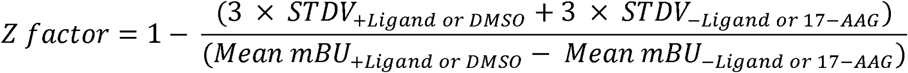

The CV value of the DMSO or 17-AAG controls was calculated as follows:

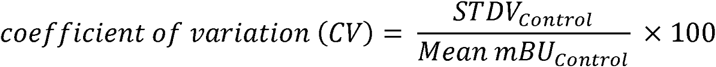

The inter-assay CV of the controls across different plates was calculated using the same equation, with the mean and STDV of the mean mBU ratio of the controls on each plate.

In the compound library screen, raw Donor and Acceptor signals were used to plot the X-Y scatter plot to visualise the separation of the positive 17-AAG and negative DMSO controls. When necessary, Donor and Acceptor signals were normalised based on the averaged values of DMSO control on each plate, which were then used to plot the X-Y scatter plot. The % inhibition of the samples was calculated based on the average mBU ratio of DMSO and 17-AAG controls on each plate using the following equation:

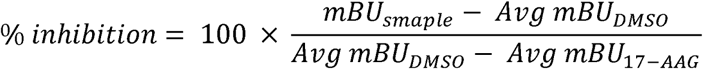

The data was analysed using Dotmatics Scientific R&D Platform (Insightful Science, LLC), where hits were identified as compounds with a % inhibition that is over or at the mean plus one standard deviation (SD). Pan assay interferers were excluded from the resulting list of hits based on two criteria; 1) the chemical structure, and 2) the raw Donor and Acceptor signal, with compounds excluded when the raw signal exceeds minus three SD.

Ordinary one-way ANOVA with Dunnett’s multiple comparisons test were used for comparisons of more than two groups and unpaired t-test was used for comparing two independent groups. Results with *p* values < 0.05 were considered statistically significant. Data are reported as mean ± SEM. The statistics and graphs were performed in Excel (Microsoft) and GraphPad Prism (GraphPad Prism 9 software).

### Cell lysate preparation

HEK293T cells transfected with HaloTag/Nluc constructed from the optimisation assay were lysed in the appropriate amount of RIPA buffer (50 mM Tris pH8.0, 150 mM NaCl, 1% TX-100, 0.5% Sodium deoxycholate, 0.1% sodium dodecyl sulphate with protease inhibitor), sonicated and stored at −20°C. Protein concentration was determined using DC Protein Assay Kit (Bio-Rad). Western blot was run with 5 µg of protein lysate.

### Solubility fractionation

The fractionation for protein solubility was performed using a protocol described by Chen et al. (2016) with some minor modifications. In brief, transiently transfected and compound treated HEK293T cells were harvested in RIPA buffer, sonicated for 30 seconds and centrifuged at 12,000 rpm for 20 minutes at 4°C. After centrifugation, the supernatant was collected as the RIPA solubility fraction. The pellet, after being washed once with RIPA buffer, was then resuspended in 10% of the original lysis volume with urea buffer (7 M Urea, 2 M Thiourea, 4% CHAPs and 30 mM Tris pH8.4) and collected as the insoluble detergent-resistant fraction.

### Western blotting and quantification analysis

Protein quantification and western blotting were performed as described before (Chen et al., 2016). Five micrograms of cell lysate from the RIPA fraction and the equivalent liquid volume from the urea fraction were loaded. Western blot quantification was performed using the image analysis software, ImageJ (http://imagej.nih.gov/ij/). Integrated band intensities were normalised to that of loading control or the RIPA fraction.

### Immunofluorescence

HEK293T cells were plated on Poly-D-lysine (P6407, Sigma) coated glass coverslip overnight at 37°C. After the appropriate treatment, cells were fixed in 4% paraformaldehyde for 30 min and washed with PBS three times for 5 min. Cells were permeabilised by incubation in 0.5% Triton X-100 (Sigma) in PBS for 15 min at room temperature, followed by blocking in 1% goat serum, PBS for 1 hour at room temperature. Cells were incubated with primary antibody diluted in blocking solution overnight at 4°C. After washing in PBS three times for 5 min, cells were subsequently incubated with fluorescent secondary antibodies diluted in the blocking solution for 1 hour at room temperature. DAPI (Sigma) was then used to stain for nuclei before being mounted on coverslips using FluorSave Reagent (Calbiochem).

## Results

### Development and optimisation of a NanoBRET assay monitoring HSF1-HSP90 interaction

To identify compounds that will disrupt the HSF1:HSP90 protein-protein interaction (PPI), we used a recently developed bio-luminescence resonance energy transfer (BRET) assay, NanoBRET™ technology, which quantitatively detects protein interactions in living cells via measuring the energy transfer between two tags fused to our proteins of interest (Machleidt et al., 2015). In the presence of PPI, the bright luminescence emission can be detected after adding the Nano-Glo® Substrate, which acts as a donor allowing energy transfer to the HaloTag for fluorescence emission. The ratio of the luminescence/fluorescence signal (mBU) reflects the PPI in cells (Figure 1A). In this study, we established the HSF1:HSP90 PPI monitoration system using HEK293 cells for their robustness and ease to transfection.

**Figure 1.**
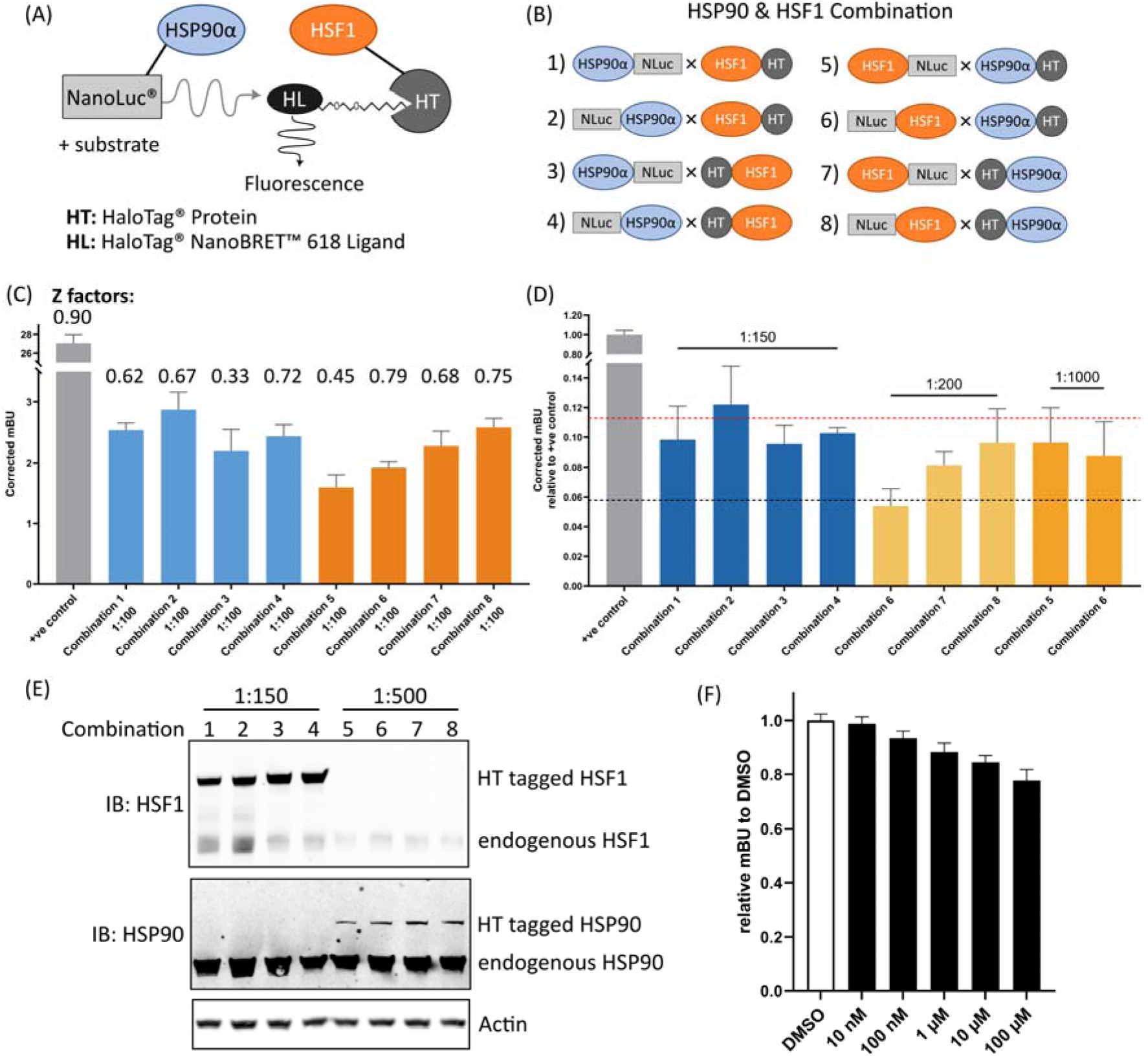
Optimising the NanoBRET system for HSF1-HSP90 interaction. **(A)** Illustration of the NanoBRET system. **(B)** Illustration of HSP90 and HSF1 combinations tested in this study. **(C-D)** Testing HSP90-HSF1 combinations and transfection ratio to maximise mBU value. HEK293T cells were transiently transfected with the testing combination pair for 48 hours before measuring the BRET signal. Blue- and orange-coloured bars represent the combination pairs with Nluc-tagged HSP90 and Nluc-tagged HSF1, respectively. The p53-MDM2 positive control is illustrated in grey colour. **(C)** Comparison of the mBU values of the eight HSP90-HSF1 combinations with the transfection ratio of Nluc donor to HT acceptor ratio as 1:100. The number indicates the Z factor between with HaloTag Ligand (HL) and without HL of each combination. **(D)** Comparison of different Nluc donor to HT acceptor ratios for HSP90-HSF1 combination. Different ratios were tested for transient transfection in HEK293T cells. The red and the black dotted line shows the maximum and minimum mBU values obtained from the Combination test with 1:100 transfection ratio shown in (C), respectively. **(E)** The expression level of the HaloTag-tagged HSP90 and HSF1. HEK293T cells were transiently transfected with Combination 1-4 and Combination 5-8 in 1:150 and 1:500 ratios, respectively. Cells were lysed after 48 hours of transfection in RIPA buffer followed by western blot analysis. Actin was used as a loading control. **(F)** The optimised NanoBRET assay shows a dose-dependent reduction of the mBU values when treated with a known HSP90 inhibitor, 17-AAG. HEK293T cells were transiently transfected with Combination 2 in 1:150 ratio for 24 hours, followed by a 24-hour treatment of different concentrations of 17-AAG. The mBU values were then determined and normalised to the mBU value obtained from cells treated with DMSO. Data represent normalised mBU values from 3 independent experiments, with mean ± SD.

To adapt the NanoBRET assay to our protein interaction of interest, both HSF1 and HSP90 proteins were tagged with either Nluc donor or HT acceptor at either the amino (N) or carboxy (C) terminus of the proteins to explore the best combination in obtaining the maximum mBU value (Figure 1B). All eight combinations of HSF1-HSP90 PPI resulted in a mBU value between 2 and 3, which is considerably lower than a p53-MDM2 positive control (Figure 1C). This is consistent with previous data showing that HSF1-HSP90 is a weak PPI and is not always detectable without chemical crosslinkers (Guo et al., 2001; Zou et al., 1998). Encouragingly, the majority of the Z factors of these combinations were above 0.5 (Figure 1C and Supplementary Figure 1A), indicating that the mBU signals detected were the robust reflection of the PPI instead of background noise. Further optimisation was carried out to adjust the Nluc donor to HT acceptor ratios. Combination 1-4, which exhibited a similar level of donor signal to the positive control (Supplementary Figure 1A), slightly enhanced the mBU value when the donor-to-acceptor plasmid ratio was further increased to 1:150 (Figure 1D). Combination 5-8, which showed a stronger initial donor signal (Supplementary Figure 1A), was tested with a harsher donor-to-acceptor ratio ranging from 1:200-1000, and those with a Z factor above zero were shown in Figure 1D. Despite the improved signal from the tested combinations, they did not exceed the performance of the earlier combinations tested (Figure 1D). The expression levels of the Nluc/HT tagged proteins from the tested combinations were also analysed by Western blot (Figure 1E). The HT-tagged HSP90 expression level was significantly lower than the endogenous HSP90 proteins, which is one of the most abundant proteins in cells and could act as a binding competitor of the HT-tagged HSP90. Strikingly, the expression level of HT-tagged HSF1 was much higher compared to the endogenous HSF1 proteins, which may be the contributing factor resulting in improved mBU values in the NanoBRET assay. As expected, the relatively low expression levels of the Nluc-tagged proteins were not detected by the Western blot. Overall, amongst the different construct combinations and transfection ratios, Combination 2 (Nluc-HSP90α × HSF1-HT) with 1:150 of Nluc donor to HT acceptor ratio we show to have the highest BRET signal (Figure 1D) and was therefore used in the following compound screen.

A known HSP90 inhibitor 17-AAG was used as a positive control to further validate this assay. 17-AAG blocks the ATPase pocket of HSP90 thereby inhibiting the function of HSP90 including its interaction with HSF1 (Jez et al., 2003; Kijima et al., 2018). Treating cells with 17-AAG resulted in a dose-dependent decrease in the mBU signal (Figure 1F). A 96-well plate assay with alternative columns of 17-AAG treatment yielded compact data points with coefficient of variation (CV) values of 1.75 and 1.58 of the treatment groups respectively of the whole plate, and an overall Z factor of 0.38 for detecting HSF1-HSP90 PPI (Supplementary Figure 2). A CV value determines the consistency of the assay, where small CV values indicate less variations of the assay. A Z factor reflects the effect size detected, with a value between 0.5 and 1 indicating a large separation while a value between 0 and 0.5 indicating a small separation (Zhang et al., 1999). Since the HSF1-HSP90 PPI is known to be a weak interaction, a Z factor above 0.2 in this study is regarded as a robust detection of the PPI. Thus, we used the cut-off point of CV value of 10 and Z factor of 0.2 in this study. As demonstrated in our optimisation, this assay produced consistent and robust readings for HSF1-HSP90 PPI.

### Screening for HSP90:HSF1 interaction disruptors

In this study, we used the 2100 Enamine Bioreference library which contains 2,100 compounds of wide varieties including FDA-approved drugs, tool compounds with validated biological activity, active metabolites and new drugs currently in clinical trials. The screening was carried out in two stages. In the first stage, we screened eight plates of library compounds including positive and negative controls (Supplementary Figure 3A). As shown in Figure 2A, the mBU values of DMSO and 17-AAG on each plate were consistent and the mBU fold change caused by 17-AAG treatment was maintained between 1.15 and 1.19 (Figure 2B). This was supported by the in-plate Z factor ranging from 0.26 – 0.78 and the CV values all below five, (Figure 2A and 2B; Supplementary Table 2), demonstrating that each plate passed the consistency and robustness threshold to identify potential HSF1-HSP90-PPI inhibitors. The assays also showed plate-to-plate consistency with an inter-assay CV of both the DMSO and 17-AAG control below five (Supplementary Table 2). The results of the complete 8 plate screening were shown in Figure 2C and D. Compounds with a % inhibition within the range of the 17-AAG control and above one standard deviation (>1SD) were identified as hits. In this stage 1 screen, Geldanamycin, a known HSP90 inhibitor, was blindly identified as one of the hits (Figure 2C and D), validating the robustness of the assay.

**Figure 2.**
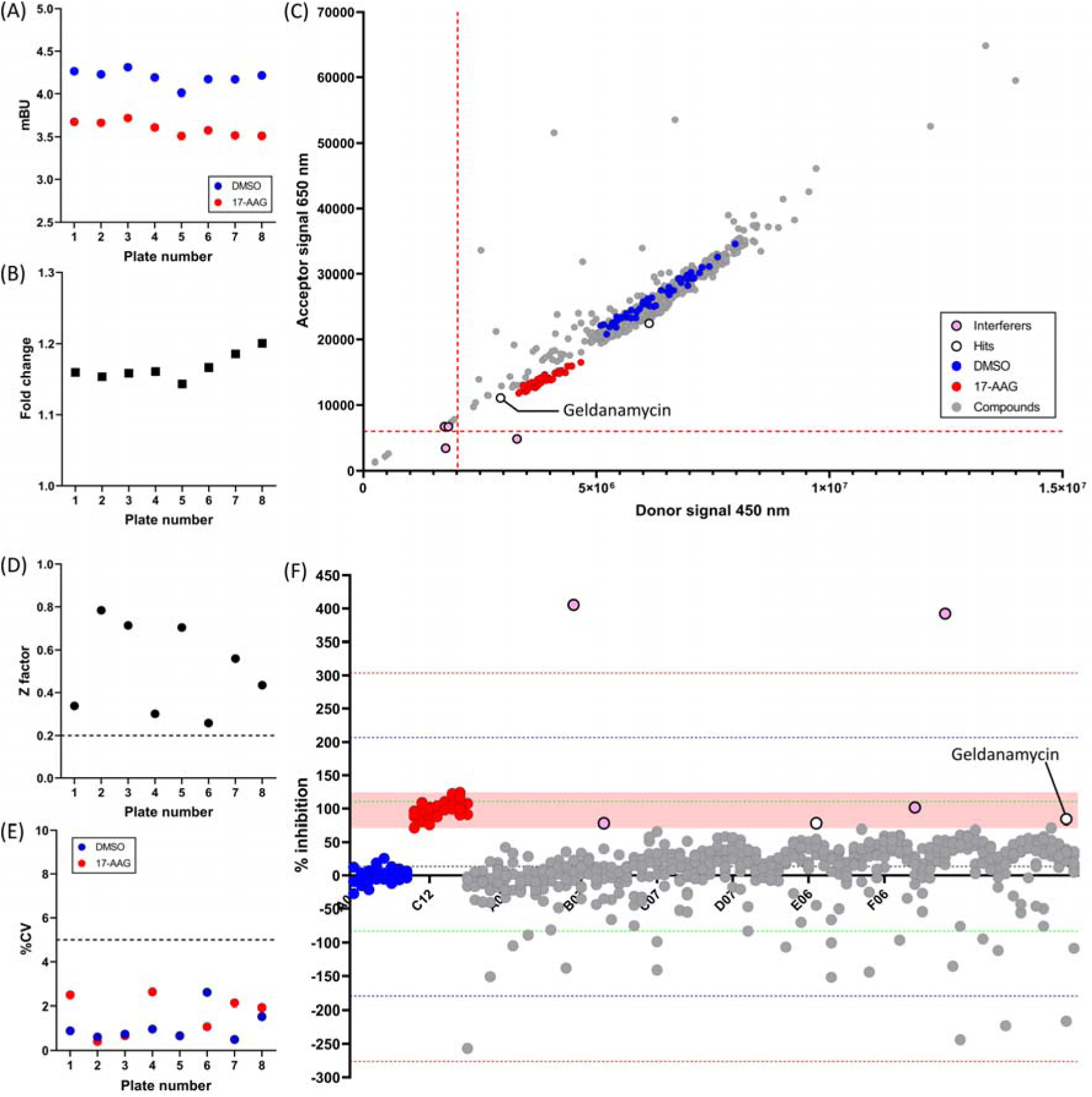
The NanoBRET assay blindly identified a known compound that disrupts the HSF1-HSP90 interaction in the first stage of 2100 Enamine Bioreference library compound screen using the NanoBRET HSF1-HSP90 interaction assay. HEK239T cells were transiently transfected with Combination 2 in 1:150 ratio for 24 hours. Transfected cells mixed with HT ligand were plated on the 96-well plate containing compounds from the Enamine Bioreference library. They were incubated at 37°C for 24 hours, after which mBU values were determined. The blue and red dots represent the DMSO and 17-AAG controls, respectively. Hits were labelled with empty black dots and interferers were labelled with pink dots. **(A-B)** Each point represents the **(A)** average mBU value of the DMSO and 17-AAG control and the **(B)** fold change between these two controls from each 96-well plate in the stage 1 screen. **(C)** Raw donor and acceptor signals from the stage 1 screen (8 plates) were plotted on a scatter plot graph. The red dotted lines represent the mean of signal minus 3SD. **(D-E)** Each point represents the **(D)** Z factor between the DMSO and 17-AAG control and the **(E)** coefficient of variation (CV) of each 96-well plate in the stage 1 screen. **(F)** The % inhibition of the compounds from the stage 1 screen (8 plates). The green, blue and red dotted lines represent the mean of % inhibition plus and minus 1, 2 and 3SD, respectively. The light red shaded region indicates the range of % inhibition of the 17-AAG controls.

Encouraged by the consistency and robustness of the assay performance, we then proceeded to stage 2 screening for the rest of the library consisting of a further 20 plates. The in-plate Z factors were all above the cut-off value (ranging from 0.43 – 0.86) and the CV values were all below five (Supplementary Figure 2D & E; Supplementary Table 4), indicating the screens were consistent and robust enough to identify PPI inhibitors. Importantly, another HSP90 inhibitor, 171009-00-0, was identified as a hit in this full screen, indicating the specificity of this assay in the second round (Figure 3A&B). Overall in the complete screen of the 2100 Enamine Bioreference library, 8 hits in total were identified (Table 1), and the ones demonstrating the highest % of inhibition were subsequently followed up (Figure 3C).

**Figure 3.**
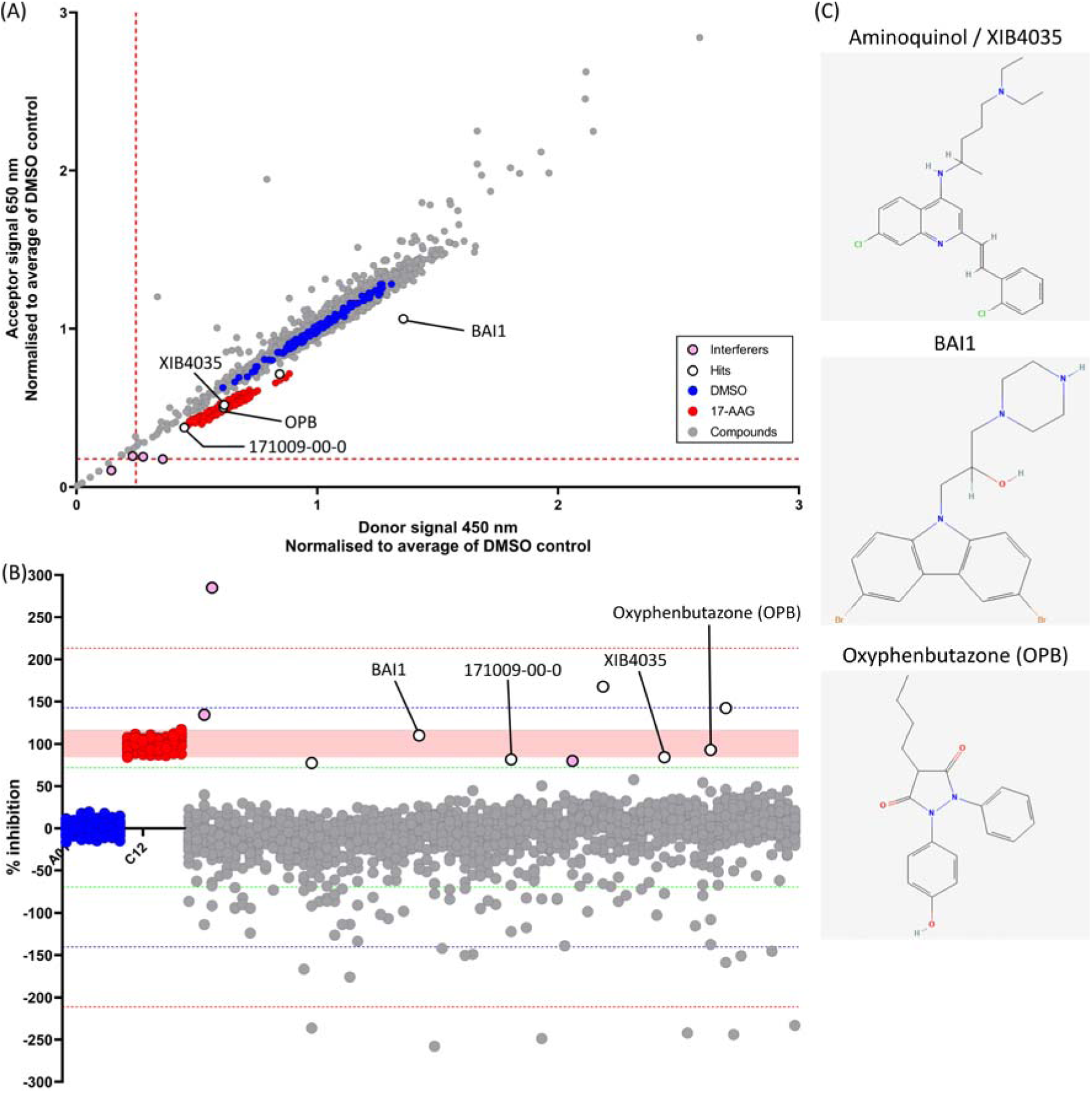
Three novel compounds were identified after the second stage of 2100 Enamine Bioreference library compound screen using the NanoBRET HSF1-HSP90 interaction assay. The rest of the 20 plates were screened using the same protocol as the stage 1 screen. The blue and red dots represent the DMSO and 17-AAG controls, respectively. Hits were labelled with empty black dots and interferers were labelled with pink dots. **(A)** Raw donor and acceptor signals from the stage 2 screen (20 plates) were normalised based on the averaged values of DMSO control on each plate. The red dotted lines represent the mean of normalised signal minus 3SD. **(B)** The % inhibition of the compounds from the stage 2 screen (20 plates). The green, blue and red dotted lines represent the mean of % inhibition plus and minus 1, 2 and 3SD, respectively. The light red shaded region indicates the range of % inhibition of the 17-AAG controls. **(C)** Compound structures of XIB4035, BAI and Oxyphenbutazone (OPB).

### OPB treatment reduces the accumulation of insoluble TDP-43 in cells

Of the three compounds, oxyphenbutazone (OPB) has previously been suggested to be a potential HSF1 activator, although its mechanism of action has not been characterised (Cervantes & Corton, 2021). The other two identified compounds were not known to associate with HSF1 previously, but BAI1 has been shown to inhibit apoptosis (Amgalan et al., 2020; Garner et al., 2019) and XIB3540, also known as Aminoquinol, has been shown to promote neurite outgrowth of Neuro-2A cells (Hedstrom et al., 2014; Tokugawa et al., 2003), which suggested that these compounds could be effective in preventing neurodegeneration.

One of the hallmarks of neurodegenerative diseases is the accumulation of aberrant protein aggregates, which causes cytotoxicity to neurons, eventually leading to neuronal death. TPD-43 protein aggregates are found in several neurodegenerative diseases, such ALS, FTD and AD (Hayes & Kalab, 2022; Meneses et al., 2021; Neumann & Mackenzie, 2019) and the presence of TDP-43 aggregates is associated with cytotoxicity both in mouse and cell models (Barmada et al., 2010; Becker et al., 2017; Chen et al., 2016; Fallini et al., 2012; Park et al., 2017; Wang et al., 2012; Yamashita et al., 2014). It is a known target of various HSPs and the activation of HSF1 is demonstrated to rescue TDP-43 aggregation and cytotoxicity (Chen et al., 2016; Lu et al., 2022; Park et al., 2017). To investigate whether the identified hit compounds can rescue aberrant proteinopathy linked to neurodegenerative disease, we tested the compound treatment in a robust TDP-43 overexpression cell model. In our initial study where TDP-43 overexpression cells were treated with 10 µM of the three tested compounds for 24 hours, despite we did not see any significant in the endogenous protein level of HSP70 and HSP90 (Figure 4A), we did observe a reduction of insoluble TDP-43 with XIB4035 and OPB treatment (Figure 4A), with a significant reduction by the OPB treatment (Figure 4B). The OPB-mediated insoluble TDP-43 clearance was dose-dependent, whereas the mode of the effect of XIB4035 was not obvious in the dosage range tested (Figure 4C and D). None of the treatment altered the level of soluble TDP-43 (Supplementary Figure 4) and we did not observe any obvious signs of cell stress or toxicity (Figure 5), indicating that the OPB reduces the level of insoluble TDP-43 potentially through facilitating protein refolding. Further study should be carried out to explore the potential of OPB as a new therapy for neurodegenerative disease.

**Figure 4.**
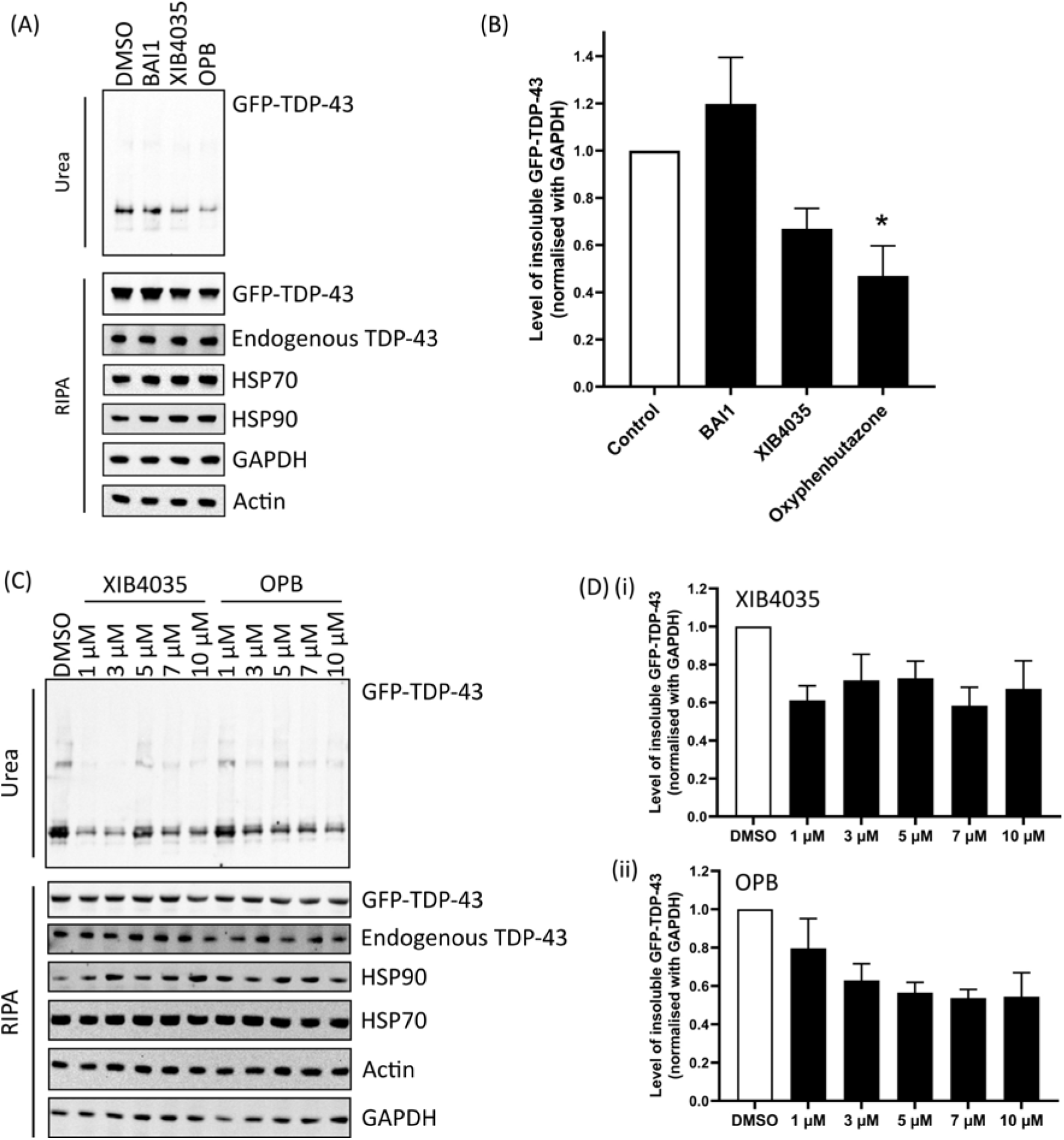
Treatment with Oxyphenbutazone (OPB) reduces the level of insoluble TDP-43 aggregates significantly. HEK293T cells were transiently transfected with GFP-TDP-43 WT for 24 hours, prior to treatment with the selected hit compounds for 24 hours at 37°C. Cells were lysed with RIPA buffer followed by fractionation. **(A)** Level of insoluble GFP-TDP-43 is detected by TDP-43 antibody in the western blot. **(B)** Levels of insoluble TDP-43 from the three independent transfections are quantified, normalised to GAPDH and shown in relative to the control, with mean ± SEM. (one-way ANOVA followed by Bonferroni post-test, *, P < 0.05). **(C)** GFP-TDP-43 WT transfected cells were treated with 1, 3, 5, 7 & 10 µM of XIB3540 or OPB for 24 hours. Level of insoluble GFP-TDP-43 is detected by a GFP antibody in the western blot. **(D)** The quantification analysis of the level of insoluble GFP-TDP-43 after 24-hour treatment of **(i)** XIB4035 and **(ii)** OPB, normalised to GAPDH and shown relative to the control, with mean ± SEM.

**Figure 5.**
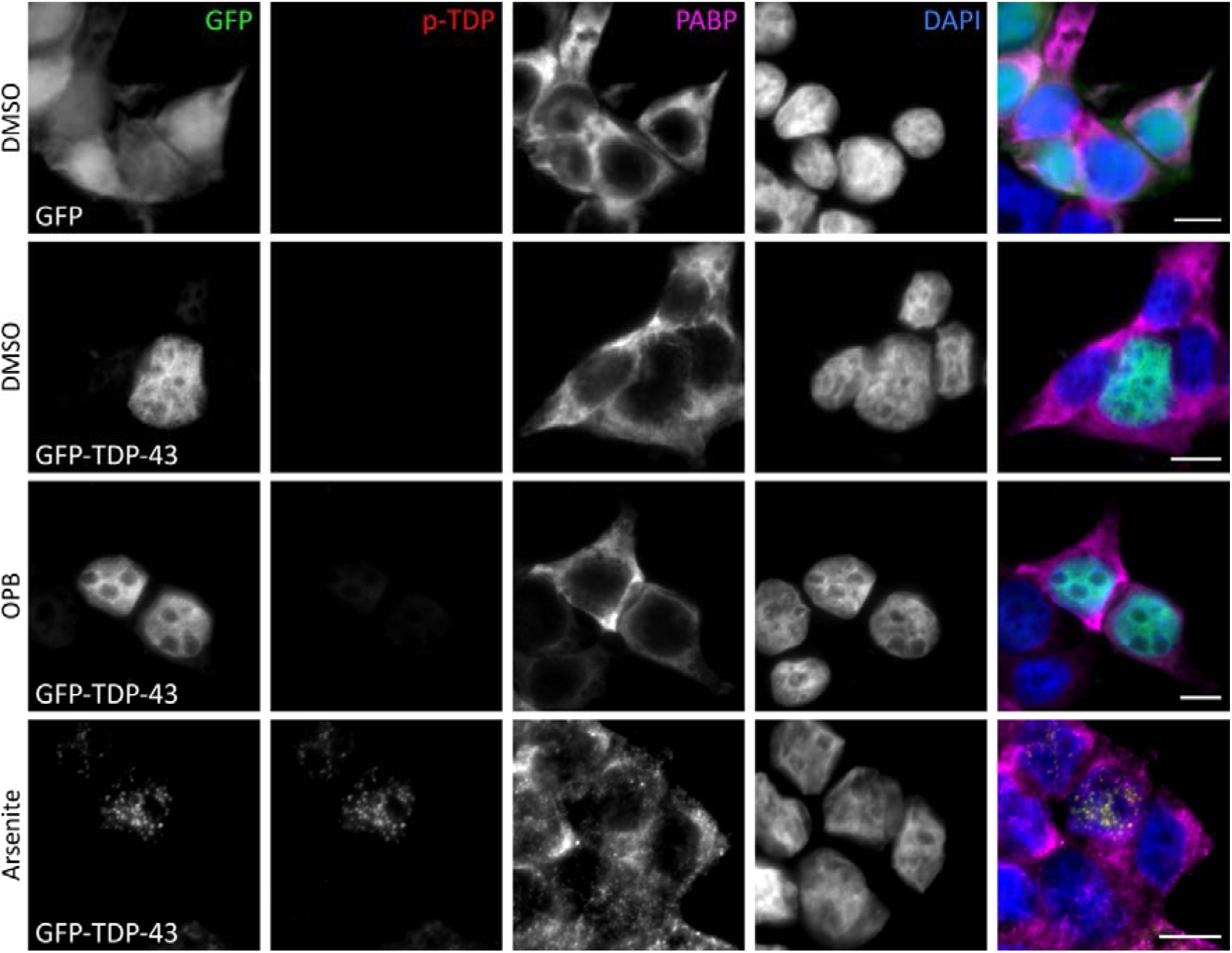
Treatment with Oxyphenbutazone (OPB) did not show any obvious signs of cell stress or toxicity. HEK293T cells were transiently transfected with GFP vector or GFP-TDP-43 WT for 24 hours, prior to treatment with DMSO or OPB for 24 hours at 37°C. Arsenite treatment (0.25 mM for 30 mins) acted as the positive control. Cells were then fixed with 4% Paraformaldehyde for 30 mins and stained for phospho TDP-43 (p-TDP) (red) and PABP (magenta). Scale bar 10 μM.

## Discussion

In this study, we established a robust screening assay to monitor the HSF1-HSP90 protein-protein interaction for drug screening in living cells (Figure 1) and identified a compound, OPB to reduce levels of insoluble TDP-43 proteins (Figure 3 and 4). OPB now serves as a potential compound for further development.

### NanoBRET assay robustly detects weak HSF1-HSP90 PPI for screening PPI inhibitors

The activation of HSF1 has long been shown to be regulated by binding to HSP90 (Zou et al., 1998). The interaction between HSF1 and HSP90 has been shown to be weak and requires a cross-linker to observe the interaction in co-immunoprecipitation assays (Guo et al., 2001; Zou et al., 1998). This interaction is linked to HSR activation that it is disrupted by heat shock or by HSP90 inhibitors like 17-AAG or Geldanamycin (Conde et al., 2009; Guo et al., 2001; Kijima et al., 2018; Zou et al., 1998). In the absence of this interaction, cells show HSF1 activation (Bharadwaj et al., 1999; Chen et al., 2014; Guo et al., 2001) and the subsequent HSPs upregulation which eventually contributes to protein aggregation clearance (Chen et al., 2016; Kundel et al., 2018; Mok et al., 2018). HSP90 is an ATP-dependent chaperone that is also involved in various cellular signalling pathways such as the PI3K/AKT and kappa-light-chain-enhancer of activated B cells (NF-κB), playing a critical role in cell survival and cytoskeleton organisation (Hoter et al., 2018). Most of the current HSP90 inhibitors block the ATPase pocket and inhibit HSP90 functions and signalling results in enhanced cell death, making inhibition of HSP90 problematic as a therapeutic intervention (Aguila et al., 2014; Garg et al., 2016; Sanchez et al., 2020). Therefore, screening for compounds that disrupt the HSP90:HSF1 interaction, but do not inhibit HSP90 function, would provide potential candidates for developing neurodegenerative disease treatment. The FRET/BRET system has been used to study protein-protein interaction for many years, used frequently in a high-throughput setting (Wu & Jiang, 2022). Here, we optimised the recently developed NanoBRET assay for this HSF1-HSP90 interaction to screen for PPI inhibitors. The assay provides only a limited signal window for compound identification with a fold change under 1.5, as shown in the optimisation assay using the known HSP90 inhibitor, 17-AAG, that disrupts HSF1-HSP90 interaction (Kijima et al., 2018) (Figure 1 and Supplementary Figure 2). This is likely partly due to the nature of the relatively weak interaction between HSP90 and HSF1 (Guo et al., 2001; Zou et al., 1998). Additionally, HSP90 is one of the most abundant proteins expressed in cells (Figure 1E; (Somogyvari et al., 2022), which could bind competitively to the HT-tagged HSF1 and deplete the interaction with the Nluc-tagged HSP90. These factors could all contribute to the low BRET signal that we observed in the NanoBRET assay.

Despite the limitations, all intra- and inter-assay CV values passed the consistency and robustness threshold (Supplementary Table 1-7) and the in-plate Z factors all met the required cut-off point (Figure 2, Supplementary Figure 3, Supplementary Table 1 and 3), indicating that the reduction of the mBU values observed reflected the disruption of the interaction. In fact, two known HSP90 inhibitors, Geldanamycin and 171009-00-0, which act similarly to the positive control 17-AAG, were identified blindly in these screens (Figure 2 and 3), demonstrating that with the appropriate optimisation and control conditions in place, this assay is robust enough to detect weak PPI and thus able to identify reliable HSF1-HSP90 PPI inhibitors with high confidence.

### OPB as a potential treatment to remove protein aggregates in neurodegenerative diseases

Among the eight compounds identified in this assay, three of the most promising on their % of PPI inhibitions in the screen and on their known bio-functional cellular effects, were followed up in this study. They have either been shown to associate with HSF1 activity such as OPB (Cervantes & Corton, 2021), or inhibit apoptosis and promote growth, such as BAI1 and XIB4035, respectively (Amgalan et al., 2020; Garner et al., 2019; Hedstrom et al., 2014; Tokugawa et al., 2003). BAI1 treatment was shown to have no or slightly positive effect in increasing the level of insoluble TDP-43 protein (Figure 4A), suggesting that it is not a promising candidate to follow up for neurodegenerative disease treatment. For XIB4035, the treatment showed a reduction in the level of insoluble TDP-43, but the decrease is not significant (Figure 4A). Furthermore, the impact on TDP-43 is not in a liner dose-dependent manner (Figure 4B) which suggests that the reduction of insoluble TDP-43 seen in XIB4035 treatment is likely to be indirect. Further study is required to characterise its mode of action.

Of the three compounds identified, OPB treatment not only generated a significant decrease in the level of insoluble TDP-43 but also acted in a dose-dependent manner (Figure 4). Despite the hints of OPB manipulating HSR, it is better known as a non-steroidal anti-inflammatory drug (NSAID) targeting the phospholipase A2 (PLA2)/cyclooxygenase (COX) (Gold et al., 2012; Saleem et al., 2018; Singh et al., 2004). Interestingly, COX enzymes have also been shown to be involved in neurodegenerative disease, where the expression of COX enzymes has been seen to be up-regulated in brain tissue of AD, PD and ALS patients (Moussa & Dayoub, 2023; Nango et al., 2023) and can be target disease treatment. For example, COX-2 specific NSAIDs such as Rofecoxib, has been shown to inhibit the COX-2 proinflammatory signalling cascades in SOD1^G93A^ mice, significantly decreasing the density of inflammatory cells and helping restore the number of motor neurons in SOD1^G93A^ mice (Moussa & Dayoub, 2023; Nango et al., 2023; Zou et al., 2020). Some NSAIDs, such as ibuprofen, indomethacin and flurbiprofen, are also involved in protein aggregates clearance by targeting the activity of γ-secretase and transcription factors, such as nuclear factor NF-κB and peroxisome proliferator-activated receptor gamma (PPARγ), to reduce the accumulation of amyloid-β protein (Moussa & Dayoub, 2023; Sastre et al., 2008). Furthermore, the inflammation pathway can also be regulated by HSPs via COX-2 and NF-κB (Kalmar et al., 2014; Schroeder et al., 2024; Wang et al., 2014) where up-regulation of HSPs reduces COX-2 expression and the production of various inflammatory mediators and pro-inflammatory cytokines in cells and *in vivo* (Ialenti et al., 2005; Thakur & Nehru, 2015). Overall, these suggest that OPB could activate HSR to refold the protein aggregates, and at the same time inhibit the inflammatory pathway *in vivo*, which eventually prevents neurodegeneration. Further study will be required to validate and explore the mechanism and effect of OPB in an ALS-relevant TDP-43 in vivo model, which will provide critical insights of OPB’s therapeutic potential for ALS.

In summary, we established a NanoBRET system for compound screening, which allows us to monitor and quantify HSP90-HSF1 interaction in living cells in a high-throughput setting. One of the compound hits, OPB, showed an effect in reducing the insoluble TDP-43, which is the hallmark aggregates found in about 95% of ALS patients. Therefore, we demonstrate that the NanoBRET assay is a useful tool for compound screening and OPB is a promising lead to develop potential new treatments for ALS.

## Funding and Acknowledgements

This research was funded principally by the Motor Neurone Disease Association (Chen/Apr17/858-791) and impact acceleration grant by the Medical Research Council to HJC, with additional support from the Alzheimer’s Research UK grant (ARUK-2021 DDI-UCL) to the Drug Discovery Institute.

