## supplementary for "Identification of novel small molecule chaperone activators for neurodegenerative disease treatment"

### **Supplementary Table 1)** Average mBU values, standard deviation (SD), coefficient of variation (CV) and Z’ factor between DMSO & 17-AAG controls of each plate in the stage 1 screen. The CV values were ranked as good (between zero – 10), acceptable (between 10 – 15) and unacceptable (above 15), which were highlighted in green, yellow, and red colour, respectively. The Z factor was ranked as good (between 0.2 – 1), acceptable (between 0 – 0.2) and unacceptable (below 0), which were highlighted in green, yellow, and red colour, respectively.

|  | **Average mBU** | | **SD** | | **Intra-assay CV** | | **Z factor** |
| --- | --- | --- | --- | --- | --- | --- | --- |
|  | **DMSO** | **17-AAG** | **DMSO** | **17-AAG** | **DMSO** | **17-AAG** |  |
| **Plate 1** | 4.2631796 | 3.6755379 | 0.0380305 | 0.0918004 | 0.892068 | 2.497604 | 0.337194 |
| **Plate 2** | 4.2273143 | 3.6642849 | 0.025752 | 0.0148473 | 1.493323 | 1.404809 | 0.783674 |
| **Plate 3** | 4.3117748 | 3.7209716 | 0.0319191 | 0.0247707 | 1.907639 | 2.535157 | 0.712139 |
| **Plate 4** | 4.1907632 | 3.6094086 | 0.040686 | 0.0949003 | 1.993567 | 2.629249 | 0.300325 |
| **Plate 5** | 4.0139861 | 3.5106103 | 0.0267325 | 0.0231092 | 2.022337 | 1.258794 | 0.702955 |
| **Plate 6** | 4.1706669 | 3.5752599 | 0.1090968 | 0.038518 | 2.615812 | 2.041626 | 0.256232 |
| **Plate 7** | 4.1701606 | 3.5173764 | 0.0206943 | 0.0751735 | 1.793547 | 2.137202 | 0.559421 |
| **Plate 8** | 4.2151576 | 3.5112687 | 0.0648025 | 0.067784 | 1.537367 | 2.4745 | 0.434912 |

### **Supplementary Table 2)** Average mBU values, SD and the inter-assay CV of the eight plates from the stage 1 screen (n=8).

| N=8 | **Average mBU** | **SD** | **Inter-assay CV** |
| --- | --- | --- | --- |
| **DMSO** | 4.195375 | 0.081951 | 1.953358 |
| **17-AAG** | 3.59809 | 0.077326 | 2.149085 |

### **Supplementary Table 3)** Average mBU values, SD, intra-assay CV and Z’ factor between DMSO & 17-AAG controls of each plate in the stage 2 screen. The non-shading and grey shading indicate plates that were screened in week 1 and week 2, respectively.

|  | **Average mBU** | | **SD** | | **Intra-assay CV** | | **Z factor** |
| --- | --- | --- | --- | --- | --- | --- | --- |
|  | **DMSO** | **17-AAG** | **DMSO** | **17-AAG** | **DMSO** | **17-AAG** |  |
| **Plate 1** | 3.571611 | 3.0868211 | 0.0538336 | 0.0382764 | 1.507263 | 1.2399941 | 0.4300007 |
| **Plate 2** | 3.5245782 | 3.033073 | 0.0245638 | 0.0302426 | 0.6969301 | 0.9970944 | 0.6654779 |
| **Plate 3** | 3.5870655 | 3.1366335 | 0.0189734 | 0.0405863 | 0.5289396 | 1.2939434 | 0.6033163 |
| **Plate 4** | 3.624433 | 3.1144671 | 0.0297844 | 0.0336069 | 0.8217672 | 1.0790583 | 0.627085 |
| **Plate 5** | 3.6171078 | 3.0844761 | 0.0289919 | 0.0266912 | 0.8015211 | 0.8653384 | 0.6863703 |
| **Plate 6** | 3.6415321 | 3.0749202 | 0.0223379 | 0.0269067 | 0.613421 | 0.8750368 | 0.739268 |
| **Plate 7** | 3.491404 | 2.9877966 | 0.0259295 | 0.0360412 | 0.7426681 | 1.2062786 | 0.6308392 |
| **Plate 8** | 3.5548911 | 3.0838916 | 0.0502118 | 0.0265416 | 1.4124699 | 0.8606528 | 0.5111246 |
| **Plate 9** | 3.4873222 | 3.0318129 | 0.0456479 | 0.0309496 | 1.3089662 | 1.0208289 | 0.4955263 |
| **Plate 10** | 3.7003777 | 3.1063453 | 0.0480484 | 0.0566547 | 1.298473 | 1.8238365 | 0.4712255 |
| **Plate 11** | 3.9986619 | 3.2607843 | 0.060786 | 0.0734733 | 1.5201573 | 2.2532411 | 0.45414 |
| **Plate 12** | 4.1220092 | 3.3503204 | 0.0144858 | 0.0715633 | 0.3514267 | 2.1360129 | 0.6654774 |
| **Plate 13** | 4.181107 | 3.3565377 | 0.0643535 | 0.0287964 | 1.5391508 | 0.8579183 | 0.6610962 |
| **Plate 14** | 4.201 | 3.3418663 | 0.04169 | 0.0213754 | 0.9923837 | 0.6396251 | 0.7797824 |
| **Plate 15** | 4.287424 | 3.3775898 | 0.0545515 | 0.0649477 | 1.27236 | 1.9228998 | 0.605975 |
| **Plate 16** | 4.1322647 | 3.2788001 | 0.0726737 | 0.040178 | 1.7586891 | 1.2253876 | 0.6033168 |
| **Plate 17** | 4.1507297 | 3.4118562 | 0.0583343 | 0.0368857 | 1.4053996 | 1.0811027 | 0.6133844 |
| **Plate 18** | 4.117743 | 3.2780393 | 0.0729169 | 0.0316406 | 1.7707973 | 0.9652298 | 0.6264486 |
| **Plate 19** | 4.1447968 | 3.3272794 | 0.0992804 | 0.0217235 | 2.3953017 | 0.6528916 | 0.5559584 |
| **Plate 20** | 4.2711085 | 3.3421574 | 0.0355834 | 0.0074516 | 0.8331191 | 0.2229587 | 0.8610205 |

### **Supplementary Table 4)** Average mBU values, SD and the inter-assay CV of the 10 plates from week 1 of the stage 2 screen.

| N=10 (Week1) | **Average mBU** | **SD** | **Inter-assay CV** |
| --- | --- | --- | --- |
| **DMSO** | 3.580032 | 0.064722 | 1.807853 |
| **17-AAG** | 3.074024 | 0.042267 | 1.374987 |

### **Supplementary Table 5)** Average mBU values, SD and the inter-assay CV of the 10 plates from week 2 of the stage 2 screen.

| N=10 (Week2) | **Average mBU** | **SD** | **Inter-assay CV** |
| --- | --- | --- | --- |
| **DMSO** | 4.160684 | 0.078134 | 1.87791 |
| **17-AAG** | 3.332523 | 0.045196 | 1.356214 |

### **Supplementary Table 6)** Average mBU values, standard deviation (SD) and the inter-assay CV of the 20 plates from the stage 2 screen.

| N=20 | **Average mBU** | **SD** | **Inter-assay CV** |
| --- | --- | --- | --- |
| **DMSO** | 3.870358 | 0.299059 | 7.726902 |
| **17-AAG** | 3.203273 | 0.136455 | 4.259875 |

### **Supplementary Table 7)** Average mBU values, SD and the inter-assay CV of the 28 plates from both the stage 2 and stage 2 screens (n=28).

| N=28 | **Average mBU** | **SD** | **Inter-assay CV** |
| --- | --- | --- | --- |
| **DMSO** | 3.96322 | 0.295568 | 7.457763 |
| **17-AAG** | 3.316078 | 0.216381 | 6.525204 |

### **Supplementary Figure 1)**

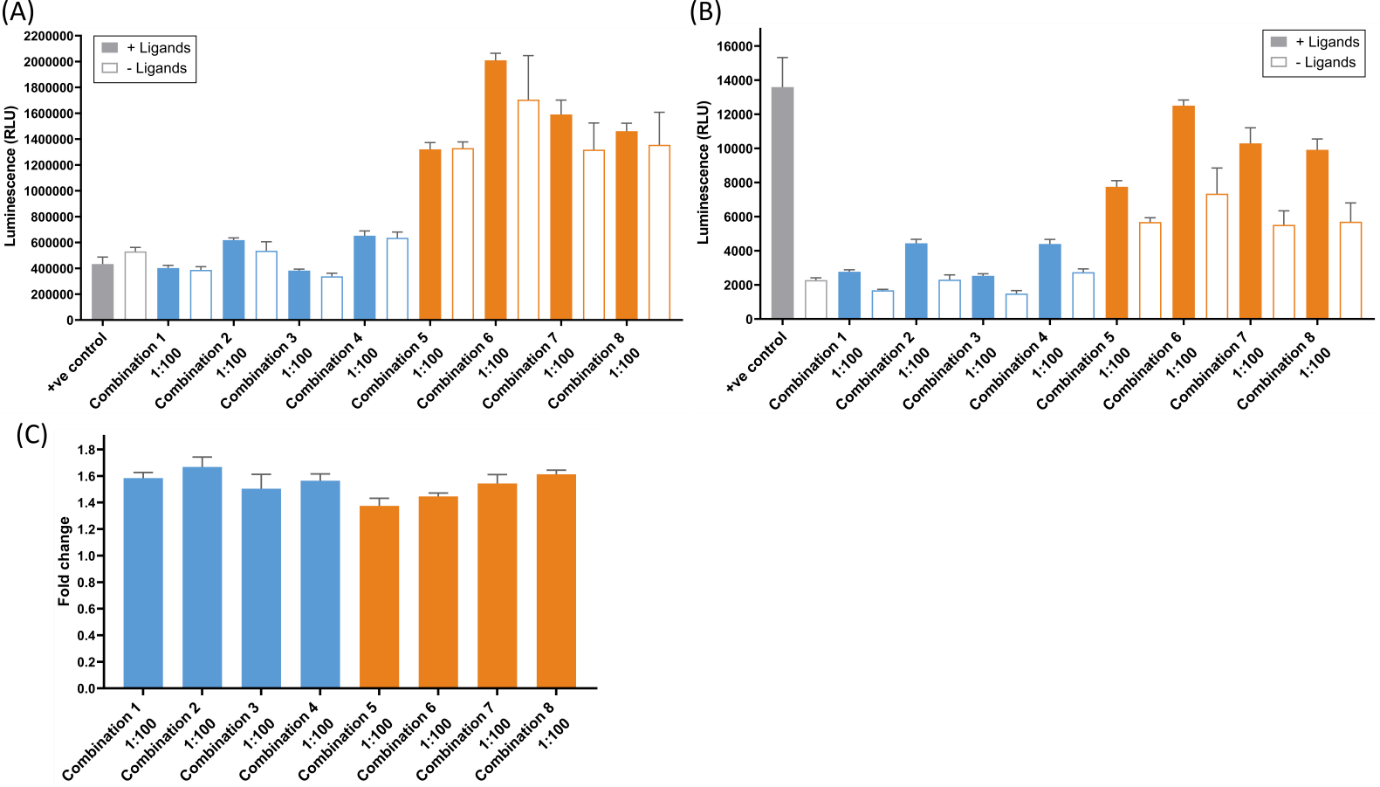

**Supplementary Figure 1)** Optimising the NanoBRET system for HSF1-HSP90 interaction. **(A-C)** Testing HSP90-HSF1 combinations and transfection ratio to maximise BRET signal. HEK293T cells were transiently transfected with the testing combination pair for 48 hours before the BRET signal was measured. The raw luminescence signal of the **(A)** Nluc donor and the **(B)** HT acceptor of the eight HSP90-HSF1 combinations and the positive control pair. The filled bars indicate the experimental pair with HT ligands and the empty bars indicate those without HT ligands. Blue- and orange-coloured bars represent the combination pairs with Nluc-tagged HSP90 and Nluc-tagged HSF1, respectively. The p53-MDM2 positive control is illustrated in grey colour. **(C)** The fold change between with and without HT ligands of the eight HSP90-HSF1 combinations, with mean values ± SD.

### **Supplementary Figure 2)**

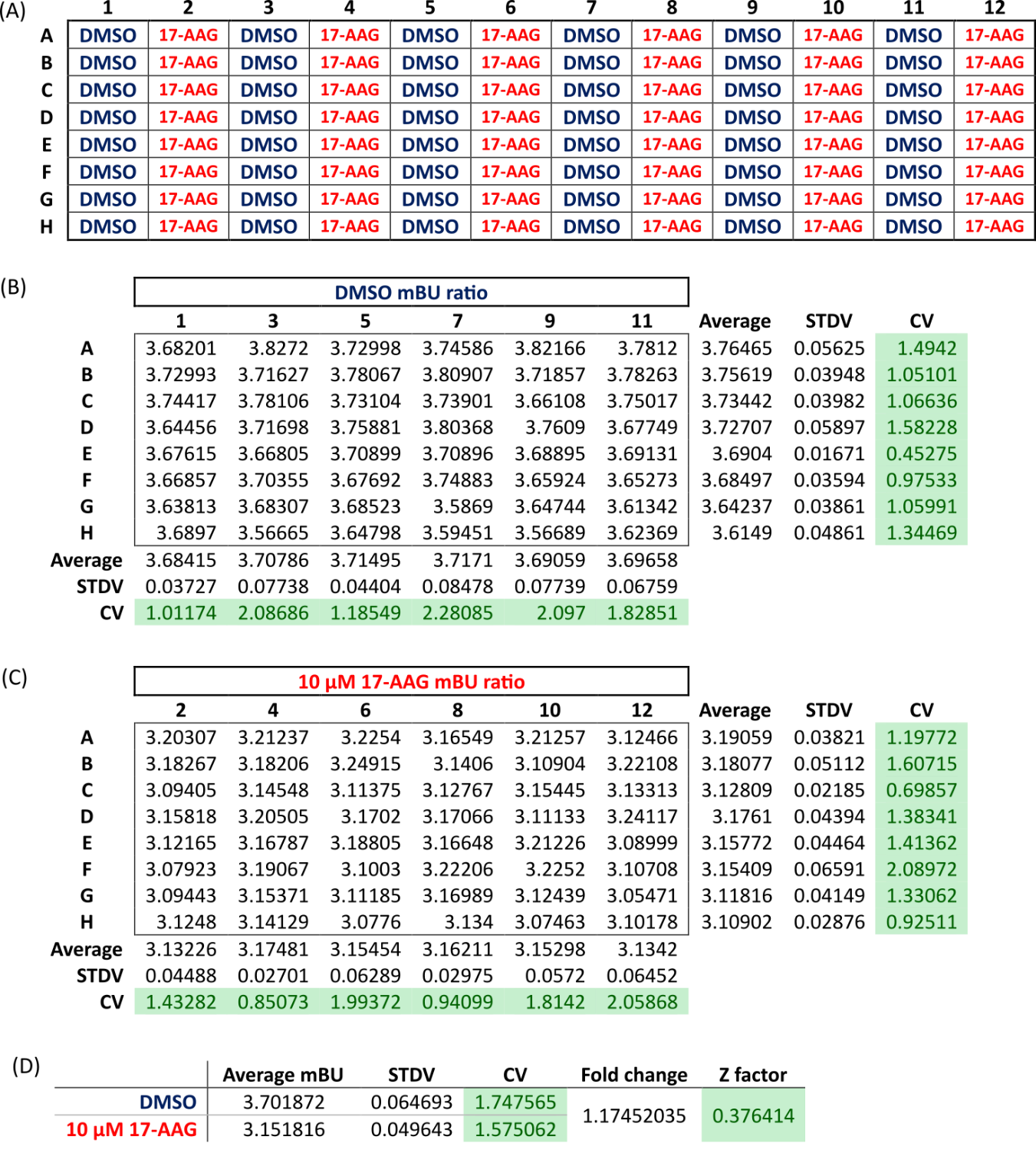

**Supplementary Figure 2)** Validation of NanoBRET HSF1-HSP90 interaction assay using a HSP90 known inhibitor, 17-AAG, with CV values and Z factor as the indicator of the feasibility and reproducibility of the assay. The CV values were ranked as good (between zero to 10), acceptable (between 10-15) and unacceptable (above 15), which were highlighted in green, yellow, and red colour, respectively. The Z factor was ranked as good (between 0.2 to 1), acceptable (between 0 to 0.2) and unacceptable (below 0), which were highlighted in green, yellow, and red colour, respectively. **(A)** The 96-well plate format for validating the NanoBRET HSF1-HSP90 interaction assay, with DMSO and 10 µM of 17-AAG controls in every other column. **(B-C)** The table shows the individual mBU value of the **(B)** DMSO and the **(C)** 17-AAG treatment using the plate format in (A). The average mBU values (Average), the standard deviation (STDV) and the coefficient of variation (CV) across the row and column were shown on the left and bottom of the table, respectively. **(D)** The table showed average mBU values, the STDV and the CV values across the whole plate of each condition. The fold change and the Z factor between the DMSO and the 17-AAG treatment were also calculated.

### **Supplementary Figure 3)**

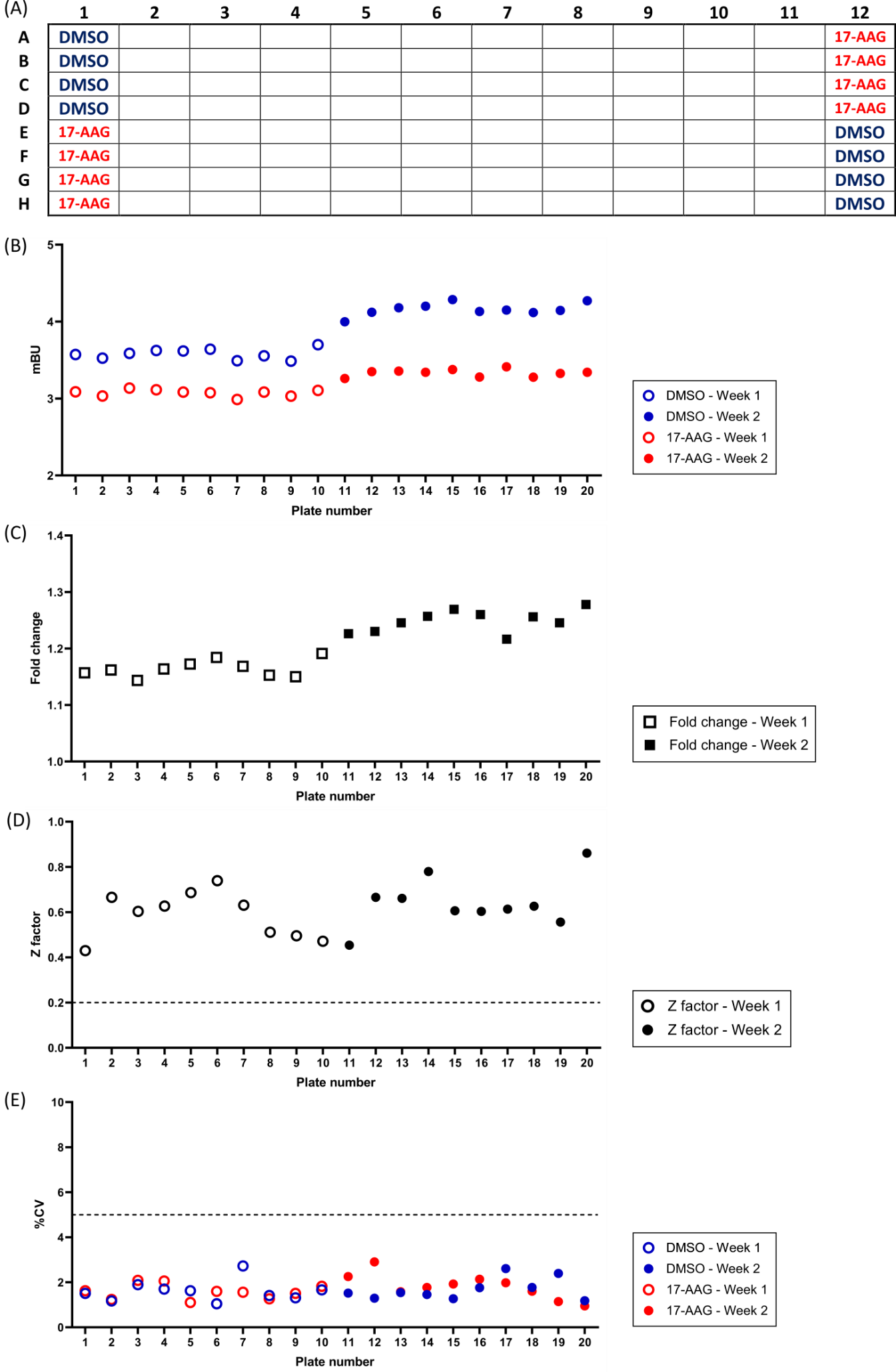

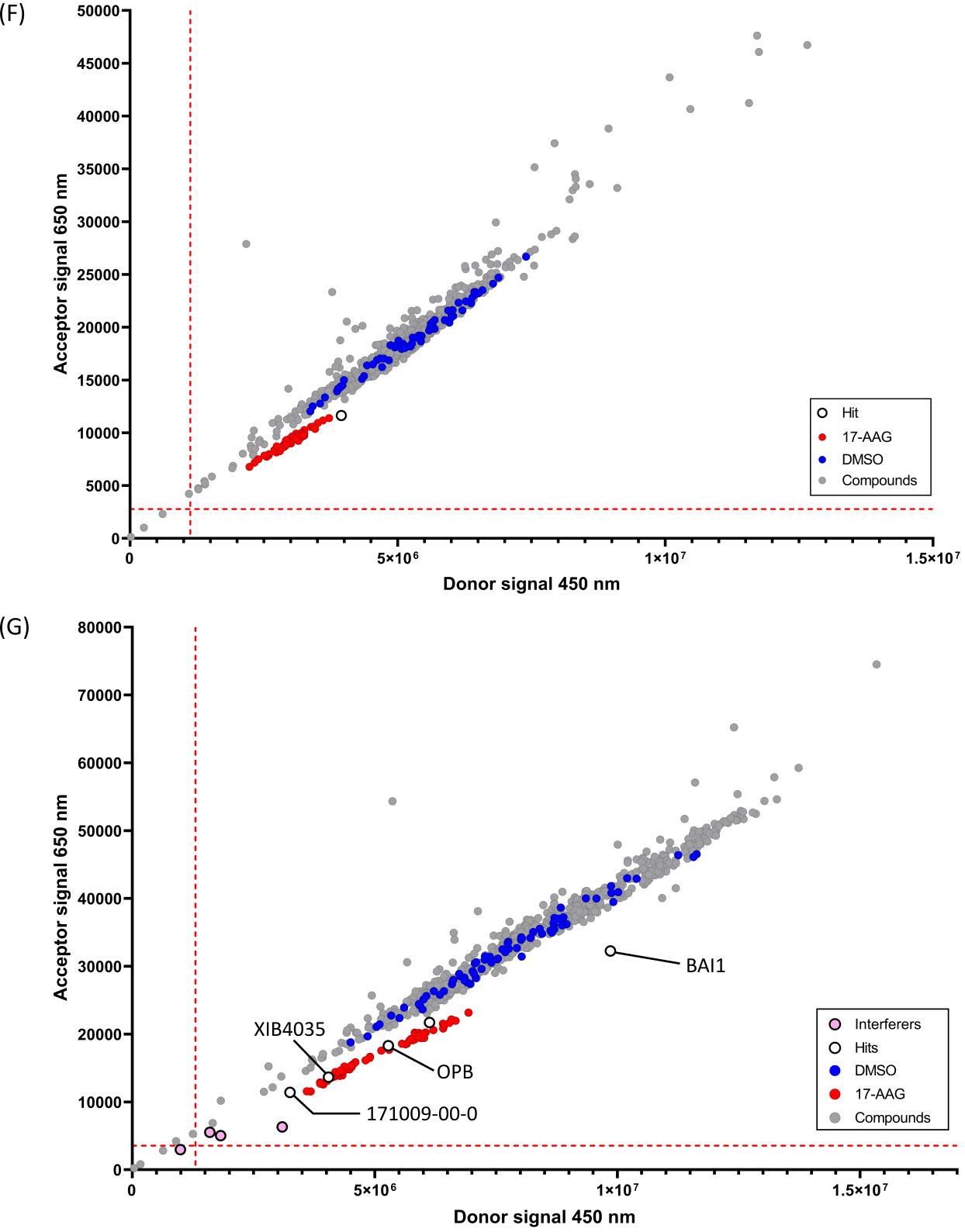

**Supplementary Figure 3)** The 2100 Enamine Bioreference library compound screen using the NanoBRET HSF1-HSP90 interaction assay. The first stage of the screen consisted of 8 plates and the rest of the 20 plates were screened using the same protocol as the stage 1 screen in two consecutive weeks (Week 1 & Week 2). The blue and red dots represent the DMSO and 17-AAG controls, respectively. Hits were labelled with empty black dots and interferers were labelled with pink dots. **(A)** The 96-well plate format for the NanoBRET HSP90-HSF1 interaction assay, with DMSO and 17-AAG controls in column 1 and column 12. **(B-E)** The DMSO and 17-AAG controls were used to indicate the feasibility and reproducibility of the assay. Each point represents the **(B)** average mBU values, **(C)** fold change, **(D)** Z factor and **(E)** coefficient of variation (CV) of each 96-well plate in the stage 2 screen. The empty and filled shapes represent data from Week 1 and Week 2, respectively. **(F-G)** Raw donor and acceptor signals from the stage 2 screen (20 plates) were plotted on two different scatter plot graphs based on the week they were screened, i.e. **(F)** Week 1 and **(G)** Week 2. The red dotted lines represent the mean of the raw signal minus 3SD.

### **Supplementary Figure 4)**

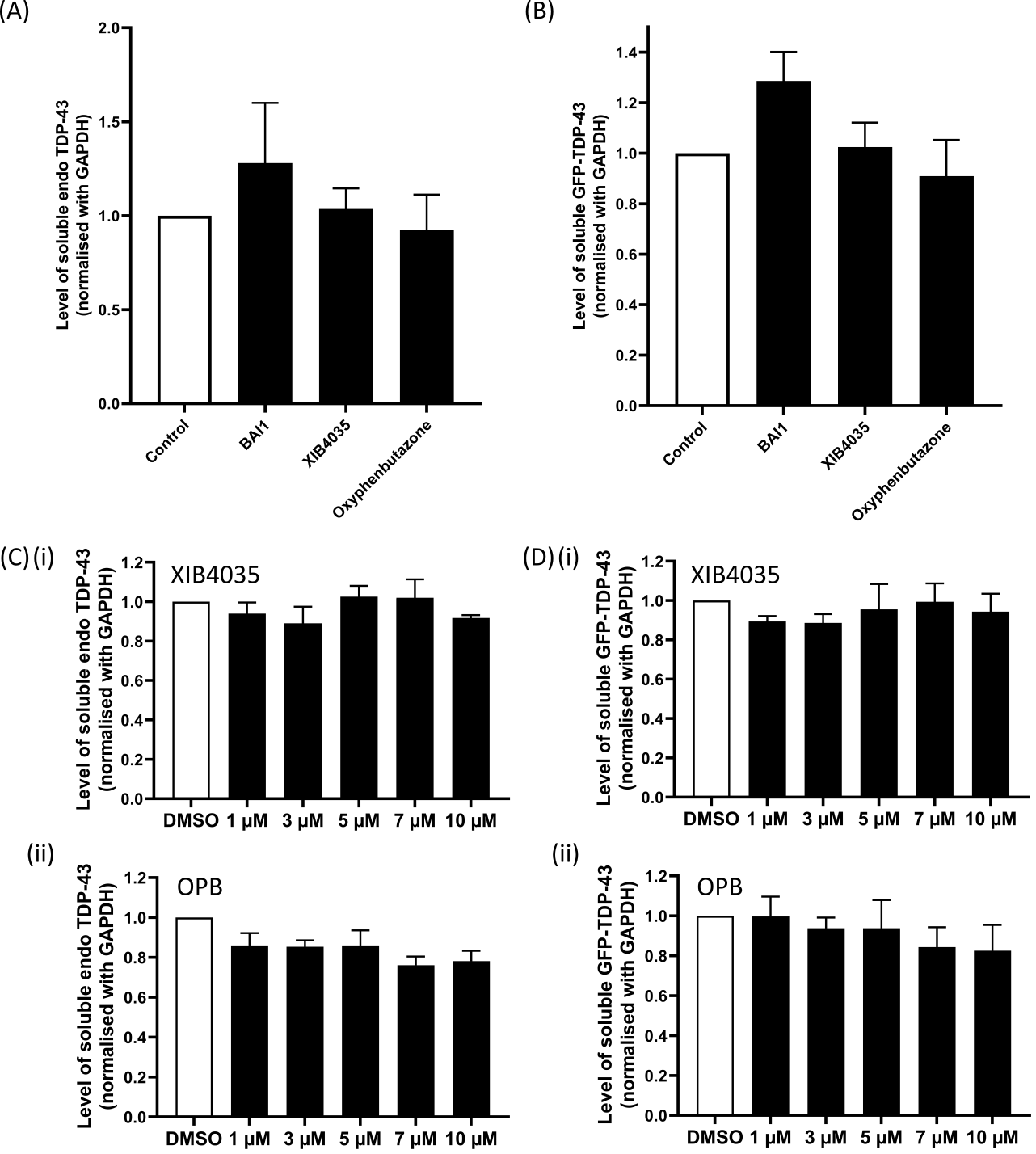

**Supplementary Figure 4)** Treatment with Oxyphenbutazone (OPB) did not affect the level of soluble TDP-43. HEK293T cells were transiently transfected with GFP-TDP-43 WT for 24 hours, prior to treatment with the selected hit compounds for 24 hours at 37°C. Cells were lysed with RIPA buffer followed by fractionation. **(A-B)** Level of insoluble GFP-TDP-43 is detected by the TDP-43 antibody in the western blot as shown in Figure 4. Levels of **(A)** soluble endogenous TDP-43 and **(B)** soluble GFP-TDP-43 from the three independent transfections are quantified, normalised to GAPDH and shown in relative to the control, with mean ± SEM. (one-way ANOVA followed by Bonferroni post-test, *, P < 0.05). **(C-D)** GFP-TDP-43 WT transfected cells were treated with 1, 3, 5, 7 & 10 µM of XIB3540 or OPB for 24 hours. Level of insoluble GFP-TDP-43 is detected by TDP-43 antibody in the western blot as shown in Figure 4. The quantification analysis of the level of **(C)** soluble endogenous TDP-43 and **(D)** soluble GFP-TDP-43 after 24-hour treatment of **(i)** XIB4035 and **(ii)** OPB, normalised to GAPDH and shown relative to the control, with mean ± SEM.
